# Seasonal dynamics of the microbial methane filter in the water column of a eutrophic coastal basin

**DOI:** 10.1101/2023.10.26.563584

**Authors:** Jessica Venetz, Olga M. Żygadłowska, Nicky Dotsios, Anna J. Wallenius, Niels A.G.M. van Helmond, Wytze K. Lenstra, Robin Klomp, Caroline P. Slomp, Mike S.M. Jetten, Annelies J. Veraart

## Abstract

In the water column of coastal waters, methane-oxidizing bacteria (MOB) can form a methane biofilter. This filter can counteract high benthic methane fluxes and thereby lower methane emissions to the atmosphere. Recent metagenomic studies revealed that the metabolism of the MOB in the filter is versatile, and could quickly respond to changing oxygen concentrations. Changes in oxygen availability in coastal basins are largely driven by seasonal stratification and mixing. However, it is still unclear how well the methane biofilter functions throughout the seasons, and how this relates to MOB community composition. Here, we determined water column methane and oxygen depth profiles and the methanotrophic community structure, methane oxidation potential, and methane fluxes of the Scharendijke basin in marine Lake Grevelingen between March and October 2021. In this period, the methane filter mainly consisted of three MOB belonging to *Methylomonadaceae*. Although in low relative abundance, the methanotrophic community was present in the mixed water column in March and had increased to 9 % by July in the stratified water column, with a distinct vertical niche partitioning in the redoxcline. The methane and oxygen gradients were vertically decoupled in summer upon the formation of a suboxic zone. Surprisingly, this did not affect the vertical distribution or potential methane oxidation of MOB. Moreover, water-air fluxes remained below 0.6 mmol m^-2^ day^-1^. Our findings suggest active methane removal by MOB in virtually anoxic water. Weakening of the stratification in September resulted in higher diffusive methane fluxes to the atmosphere (up to 1.6 mmol m^-2^ day^-1^). This was likely due to a faster supply of methane, but also a reduction of methane oxidation. Thus, despite the rapid adaptation and versatile genomic potential of the MOB community, seasonal water column dynamics significantly influence methane removal efficiency.

## Introduction

Coastal ecosystems are the major contributor to global ocean methane emissions (6–12 Tg CH_4_ yr^−1^) (Borges *et al*., 2016; Weber *et al*., 2019). Due to the presence of methane-oxidizing microorganisms in the water column and in the sediment, these methane emissions are only a fraction of the methane produced in the sediment. The efficiency of this so-called methane biofilter is one of the greatest uncertainties in methane-emission predictions (Dean *et al*., 2018). In coastal basins, a consortium of anaerobic methane-oxidizing archaea with sulfate-reducing bacteria build the methane biofilter in the sediment (Wallenius *et al*., 2021), and aerobic methane-oxidizing bacteria dominate the methane biofilter in the water column (Reeburgh, 2007; Steinle *et al*., 2017; Steinsdóttir, *et al*., 2022a; Venetz *et al*., 2023). Deoxygenation and eutrophication can lead to increased methane production in the sediment, that can exceed the methane oxidation capacity, which can result in high benthic methane fluxes into the water column (Egger *et al*., 2016; Zhang *et al*., 2020; Lenstra *et al*., 2023; Żygadłowska, *et al*., 2023a). As coastal ecosystems are especially affected by eutrophication and hypoxia (Diaz and Rosenberg, 2008; Breitburg *et al*., 2018), benthic methane fluxes are likely to increase in the future. A better understanding of the fate of methane in the water column is therefore crucial for better predictions of methane emissions to the atmosphere and key for adequate policymaking.

Microbial removal of methane in the water column is mostly attributed to putative aerobic methanotrophic bacteria, that thrive along the methane oxygen counter-gradient. Although methanotrophic archaea such as ANME-related phylotypes and NC10 bacteria can be present in the water column of marine ecosystems (Schubert *et al*., 2006; Thamdrup *et al*., 2019), aerobic gamma-or alpha-proteobacterial methanotrophic bacteria (γ- and α-MOB) dominate these methanotrophic communities (Tavormina *et al*., 2010; Steinsdóttir, *et al*., 2022b), even in hypoxic waters (Steinle *et al*., 2017). During summer stratification, such a methane biofilter can counteract the high benthic methane fluxes (Mao *et al*., 2022; Steinsdóttir, *et al*., 2022b). Multiple potential metabolic pathways within the ambient methanotrophic community ensure the functionality of the methane biofilter along the vertical range of oxygen and methane concentrations (Hernandez *et al*., 2015; Venetz *et al*., 2023). Such niche partitioning over a range of oxygen concentrations likely aids the overall resilience of the methane biofilter towards regular changes in oxygen availability in the water column(Venetz *et al*., 2023).

Coastal basins are highly influenced by seasonal water column dynamics which affects the microbial community structure and activity (Gilbert *et al*., 2012; Wang *et al*., 2020). A recent study demonstrated an effect of seasonal dynamics in MOB community and higher methane oxidation rates in summer compared to spring in bottom waters (Gründger *et al*., 2021). To reduce the uncertainties in methane-emission predictions, it is thus important to understand how the methanotrophic community structure and potential removal activity at different depths respond to the seasonal changes in methane, oxygen, and nutrient availability and how this ultimately affects methane release to the atmosphere (Louca *et al*., 2016; Grossart *et al*., 2020).

Here, we studied the seasonal dynamics of the microbial methane biofilter in the water column of a marine coastal basin in the Southwest Netherlands. In nine sampling campaigns between March and October 2021, we investigated the vertical distribution of methane and oxygen, the potential methane oxidation rates, the microbial community structure, and the methane fluxes to the atmosphere.

## Experimental procedures

### Fieldwork location and sampling methodologies

Lake Grevelingen is a highly eutrophic former estuary with a total surface area of 115 km^2^ and an average water depth of 5.1 m. The water column of the main channel is stratified during the summer months with hypoxic or anoxic bottom waters (Wetsteijn, 2011; Hagens *et al*., 2015). A more detailed description of the system and the study site can be found elsewhere (Egger *et al*., 2016; Sulu-Gambari *et al*., 2017). At the deepest point of Lake Grevelingen, the Scharendijke basin (51.742°N; 3.849°E, 45 mbs), the high sedimentation rates and anoxic bottom water lead to high benthic methane fluxes to the water column during summer stratification (0.6 - 2.7 mmol m^-2^ d^-1^) (Egger *et al*., 2016; Żygadłowska, *et al*., 2023a). To monitor seasonal water column methane dynamics in the Scharendijke basin, we conducted nine research cruises with the RV Navicula between March and October 2021. During these campaigns, we measured *in situ* water-air methane fluxes, constructed depth profiles of water column chemistry, and took samples for microbial analysis, including incubation experiments to measure potential aerobic methane oxidation rates.

The extent of water column stratification during each sampling campaign was determined with a CTD unit (SBE 911 plus, Sea-Bird Electronics, Bellevue (WA), USA). In addition, the oxygen distribution was simultaneously recorded by a seabird sensor (Seabird SBE43.) Because of the limit of detection commonly observed in oxygen sensor data of oxygen-depleted waters, we here consider anoxia once concentrations are below < 3 μmol L^-1^and do not further decrease with depth.

Water samples were taken at 30 depths with an 10 L Niskin bottle. Subsequently, unfiltered water was collected in 1 L sterile plastic bottles for DNA analysis, and 0.5 L sterile Schott bottles for incubation experiments, which were stored in the dark at 4 °C until further processing. Furthermore, 120 mL borosilicate serum bottles were filled for the determination of the methane concentration. To avoid air contamination, the bottles were filled from the bottom via gas-tight tubing while letting the water overflow three times, after which they were crimp-capped with an aluminum cap and a butyl stopper. To stop microbial activity, 0.25 mL of HgCl_2_ (sat.) was added. Samples were stored upside down at room temperature until further processing.

### Methane concentration measurements

To determine methane concentrations in the water column, 5 mL of N_2_ gas was added to all borosilicate bottles, while simultaneously removing the same volume of the water. After equilibrating for at least 2 h, methane concentrations were measured with a Thermo Finnigan Trace™ gas chromatograph equipped with a Flame Ionization Detector (detection limit: 0.02 *μ*mol L^-1^).

### DNA extraction, 16S rRNA gene sequencing, and data analysis

Water samples were filtered on Supor® PES 0.22 μm filters with a vacuum pump set up. After immediate freezing at -80 °C, the samples were stored at -20 °C until extraction. The DNA was extracted with the FastDNA™ SPIN Kit for Soil DNA isolation kit (MP Biomedicals) according to the protocol.

The V3-V4 region of the 16S rRNA gene was sequenced (Illumina MiSeq platform, Macrogen, Amsterdam, the Netherlands) to analyze the microbial community composition. The primer pairs Bac341F (CCTACGGGNGGCWGCAG) (Herlemann *et al*., 2011) and Bac806R (GGACTACHVGGGTWTCTAAT) (Caporaso *et al*., 2012) were used for bacteria and the primer pairs Arch349F (GYGCASCAGKCGMGAAW) (Takai Ken and Horikoshi Koki, 2000) Arch806R (GGACTACVSGGGTATCTAAT) (Takai Ken and Horikoshi Koki, 2000) (Takai Ken and Horikoshi Koki, 2000) were used for archaeal 16S sequencing. Seqencing data was processed with RStudio. Primers were removed with *cutadapt* (Martin and Rahmann, 2012) with the options -g, -G, and --discard-untrimmed. Low-quality reads (< Q20 forward and < Q30 reverse) were removed by truncating reads to a length of nt 270 forward nt 240 reverse with the DADA2 pipeline (Callahan et al., 2017). Finally, amplicon sequence variants (ASVs) were inferred, forward and reverse reads were merged, and chimaeras removed. For taxonomic assignment, the 254 Silva non-redundant train set v138 (https://zenodo.org/record/3731176#.XoV8D4gzZaQ) was used. For further clustering and calculation of relative abundances the *Phyloseq* package was used, and data was visualized with ggplot2, and graphs were adjusted with Adobe Illustrator. Raw reads of the 16S amplicon sequencing data can be accessed on the European Nucleotide Archive under the accession number (TBD).

### Potential methane oxidation rates

To investigate the seasonal dynamics of aerobic methane removal potential by the methanotrophic methane filter in the water column, we incubated water samples from 14 depths during each sampling campaign within 24 h after sample retrieval. For each depth 100 ml of unfiltered, air-equilibrated sample, was put into an autoclaved 120 mL borosilicate bottle (in triplicate) and closed with bromo-butyl stoppers, and crimped with an aluminum cap. To each incubation, 1 mL of ^13^C-CH_4_ (99%) was added, which resulted in a partial pressure of 5 % in the headspace. All bottles were incubated under constant shaking (150 rpm) in the dark, at room temperature for the duration of the incubation.

For each time point, a 1 mL liquid sample was taken from the incubation bottle and replaced with 1 ml of air. As the carbonate balance at the pH range of our sample is susceptible to small changes in pH, we acidified subsamples to a pH of < 2. The liquid sample was transferred into a gas-tight air equilibrated 3 mL vial (Labco, exetainer, UK) containing 50 μL of 0.1 M HCl. The produced ^13^C-CO_2_ was measured directly from the exetainer headspace with a GC-MS (Agilent 5975C inert MSD). Liquid ^13^CO_2_ concentrations were calculated with the Henry coefficient (Supplementary Infomration). The linear increase in ^13^C-CO_2_ after the lag phase was used to determine methane oxidation rates.

### In situ water-air flux measurements

*In situ* water-air fluxes of methane were measured during each sampling campaign. Fluxes were determined using a transparent, cylindrical floating chamber (ø: 390 cm, height 27 cm, TechnoCentrum, Radboud University, Nijmegen, NL) connected in a closed loop to a LICOR trace gas analyzer (LI-7810, LI-COR Environmental - UK Ltd, Cambridge, UK). The chamber frame was stabilized to withstand wave turbulence using a bespoke raft (Supplementary Information Figure S1), as the surface water of the Scharendijke basin can be quite turbulent. To ensure a closed loop between the chamber and the LICOR gas analyzer, the input and output connectors of the LICOR were connected to the top of the floating chamber using gas tight polyurethane tubes (ø: 4 mm (inside), Festo, 5 m). The chamber was gently placed on the water surface and the accumulation of methane was measured with the trace gas analyzer for at least three minutes, in triplicate. The chamber was aerated until atmospheric methane concentrations were reached, before starting each new measurement.

Methane fluxes were then calculated with the following equation:

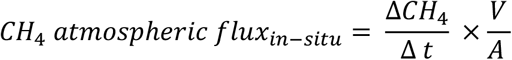

−where ΔCH_4_ /Δt is the linear increase of the concentration of methane (mmol m^-3^) in the chamber over time (Δt). V is the volume of the chamber (m^3^) and A is the area of the chamber (m^2^). The measured partial pressures (ppb) in the chamber were converted to methane (mmol m^-3^) using the ideal gas law and the ambient air temperature during each deployment.

### Statistical analysis

To assess the seasonal dynamics of the bacterial community, Shannon Diversity and Chao1 richness were calculated to illustrate the bacterial alpha diversity of untransformed ASV counts of each sample. Before analysis, ASVs that were assigned to archaea, mitochondria and chloroplasts were removed from the dataset.

The dissimilarity of the bacterial community between each sample was calculated as the Bray-Curtis distance of rarefied ASV counts and distance matrices were illustrated by non-metric multidimensional scaling (NMDS). Based on the stress factor assessment of the ordination with different dimensions, the ordination was performed on two dimensions (stress = 0.092, nonmetric fit R^2^ = 0.992 and linear fit R^2^ = 0.965). Vectors for the environmental variables O_2_, CH_4_, H_2_S, NH_4_, NO_2_, NO_3_, depth were determined with the *env_fit()* function of the *vegan* package.

## Results and discussion

In this study, we monitored the seasonal dynamics of the microbial methane filter in the water column of a marine basin during nine sampling campaigns in 2021. In the following sections, we will show and discuss our results for the seasonal succession and distribution of MOB, methane oxidation potential along the depth gradient, the potential for anoxic methane removal, and the water-to-air fluxes of methane in marine Lake Grevelingen in 2021. We show that there is 1) a vertical distribution and seasonal succession of the versatile methanotrophic community, 2) that this is related to water column dynamics and the availability of oxygen and nutrients and 3) that this is ultimately related to methane removal and *in situ* methane fluxes to the atmosphere.

### Seasonal succession and vertical distribution of the methanotrophic community

The methanotrophic community was dominated by methane-oxidizing Gammaproteobacteria (γ-MOB) belonging to *Methylomonadaceae*. Although Alphaproteobacteria (α-MOB) were present, their relative abundance never exceeded 0.07 % of the bacterial 16S rRNA reads. This could be due to the eutrophic state of the basin, as α-MOB are more adapted to oligotrophic conditions and could be continuously outcompeted by *Methylomonadaceae* (Ho *et al*., 2013; Kaupper *et al*., 2020). The main methanotrophic genera were *Milano-WF-1B (M1B), Marine_Methylotrophic_Group2* (*MMG2*), and *Methyloprofundus (MP)*, which together built a methanotrophic community that is potentially versatile in terms of oxygen metabolism, denitrification capacity and potential for sulfur transformation (Venetz et al. 2023). Despite the low diversity of the methanotrophic community, our seasonal data demonstrated a high adaptation potential enabling both vertical niche partitioning and seasonal succession. While the relative abundance of MOB was less than 0.3 % of the bacterial 16S rRNA reads in the mixed, oxygenated water column in March (Figure 1A), it increased up to 9 % at 39 m depth by July when the water column was stratified (Figure 2A). This drastic increase can be attributed to the summer stratification which resulted in the formation of chemical niches along the methane-oxygen counter gradient (Amaral and Knowles, 1995; Mayr *et al*., 2020) and was accompanied by a shift in the MOB community structure with depth: *M1B* became dominant in the oxygenated water but was outcompeted by *MMG2* and *MP* in the anoxic waters (Figure 2A). A similar niche partitioning was observed at the end of summer stratification in September 2020 (Venetz *et al*., 2023). There, the metabolic potential of these genera indicated metabolic adaptations to oxygen limitation with high affinity oxidases and through potential nitrate, iron and sulfur reduction, which could explain the shift in community composition along the oxycline. Similarly, these metabolic adaptations are also important for seasonal succession. Our seasonal data showed that despite having low diversity, the methanotrophic community could adapt to both water column mixing and summer stratification. However, eutrophication and stratification in coastal ecosystems will likely intensify in the future (Dominović *et al*., 2023), which may induce more pronounced shifts in the methanotrophic community during summer stratification and further decrease diversity. The diversity of the entire water column bacterial community indeed shows a drastic decrease in alpha diversity from March to October in 2021 (Figure 2C). Prolonged anoxia and warming could arguably promote even slow-growing methanotrophic archaea or NC10 bacteria (Su *et al*., 2023). Yet, the pool of other methanotrophic microorganisms than *Methylomonadaceae* in marine Lake Grevelingen is very low. Thus, the low diversity of methanotrophic microorganisms can impair the resilience towards changes in oxygen availability that exceed the amplitude of the current seasonal dynamics.

**Figure 1:**
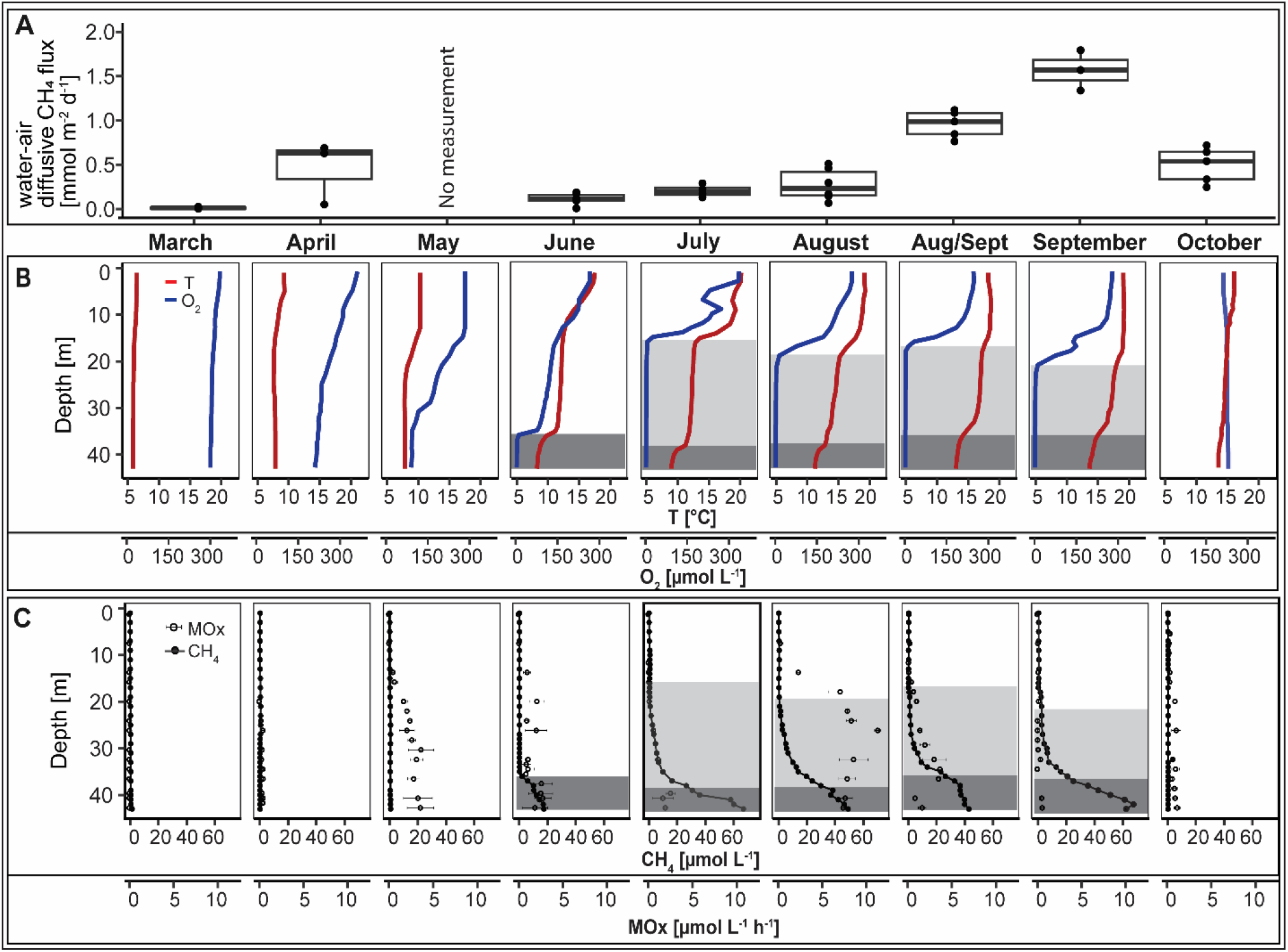
**A** Diffusive methane water-air fluxes measured with an in situ floating chamber. Boxes indicate the first and third quartiles, lines indicate the median, whiskers indicate outer data points if less than 1.5 interquartile range from quartiles. **B** Depth profiles of oxygen (blue lines) and temperature (red lines), and **C** methane concentrations (full circles) together with potential aerobic methane oxidation rates (MOx) (empty circles) determined by incubation experiments between March and October 2021 in the Grevelingen Scharendijke basin. Error bars of MOx rates show standard deviation between the biological replicates. The hypoxic zone (< 63 μmol L^-1^ (Breitenburg et al., 2018)) is indicated in light grey and the anoxic zone (< 3 μmol L^-1^) is indicated in dark grey.

**Figure 2:**
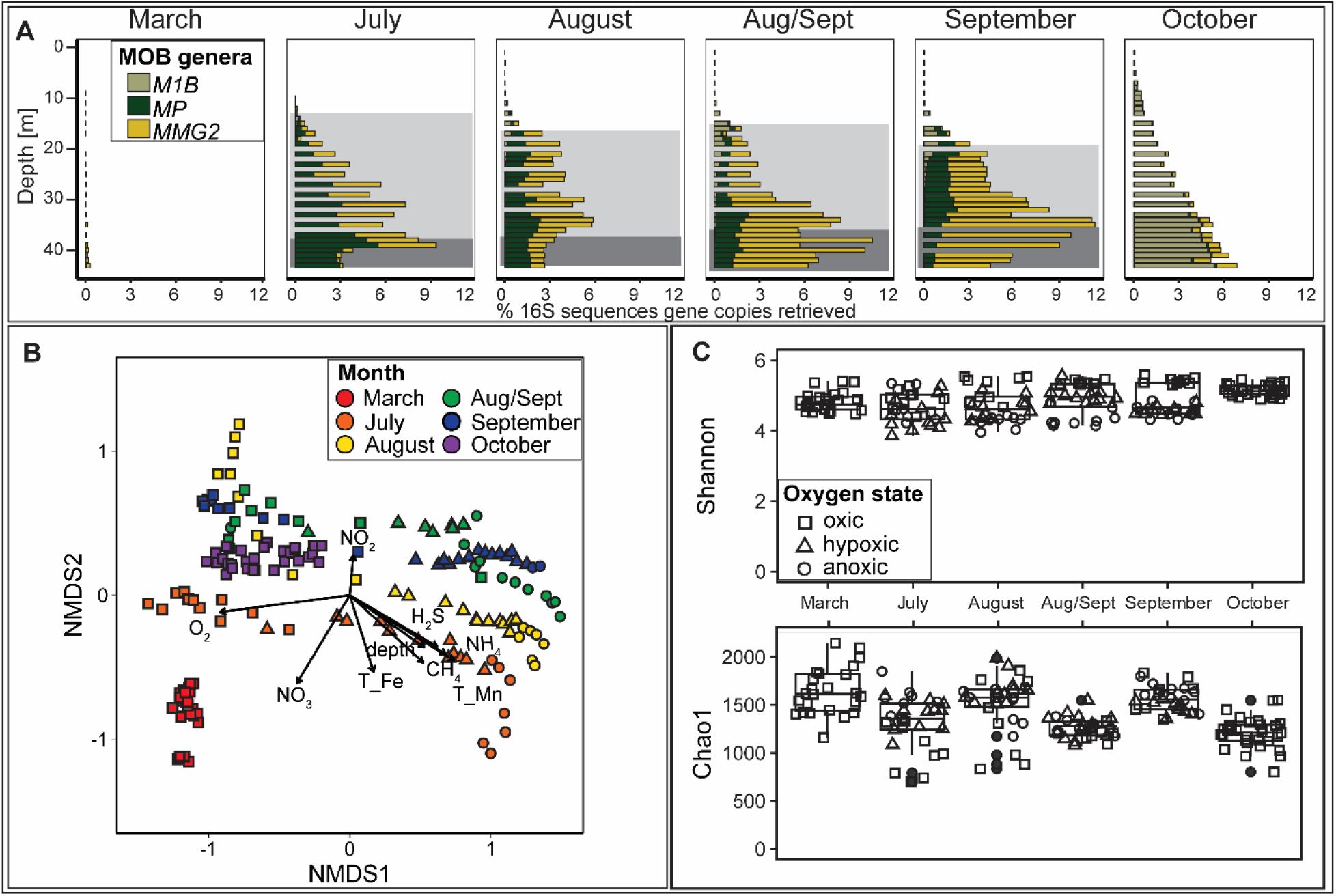
**A** Relative abundance of methanotrophic bacteria retrieved through 16S rRNA sequencing. Light grey areas indicate the hypoxic zone (< 63 μmol L^-1^ h^-1^) and dark grey areas indicate the anoxic zone (< 3 μmol L^-1^). **B** Alpha diversity measures Chao1 (richness) and Shannon (richness and evenness) of the total water column bacterial community of each month. **C** Beta diversity of all samples calculated as Bray-Curtis distance and ordinated via two-dimension NMDS. Short arrows indicate weak correlation and long arrows indicate strong correlations. The R^2^ values can be found in the Supplementary Data (“NMDS_output”).

### Methane removal by methanotrophic bacteria

Potential aerobic methane oxidation rate measurements (up to 0.60 μmol L^-1^ h^-1^) indicate that benthic methane is (partially) oxidized in the water column before reaching the atmosphere (Figure 1C). These measurements confirmed that there is potential for methane removal throughout the year. Moreover, methane oxidation rates followed a seasonal pattern, and varied with depth along the water column. Methane oxidation in the mixed water column in March was low (< 0.002 μmol L^-1^ h^-1^) compared to May (0.008 – 3.6 μmol L^-1^ h^-1^) upon the onset of summer water column stratification and the development of an oxycline. During the stratification period, methane oxidation rates followed a vertical pattern. In June a clear methane-oxygen counter gradient had formed with an oxycline between 35 and 36 m and high methane concentrations in the bottom water (up to 17 μmol L^-1^). The high methane oxidation rates below 9 m (0.8 - 2.9 μmol L^-1^ h^-1^), indicated active aerobic methane removal. However, canonical aerobic methane oxidation alone cannot explain the observed biogeochemical and microbiological pattern, which is especially pronounced in July, August and September. Between June and July, an additional thermocline formed in between 12 and 16 m (Figure 1B). This induced the formation of the oxycline higher up and resulted in a vertical decoupling of the methane and oxygen gradients and the formation of an oxygen- and sulfide free (suboxic) zone (Żygadłowska, *et al*., 2023b). Interestingly, the potential aerobic methane oxidation rates and the relative abundance of MOB were highest far below the oxycline (Figure 1 C: Figure 2A). If this de-coupling had induced a diminishing of the methane biofilter, we would have expected a decreased abundance of MOB, lower methane oxidation rates and a higher methane flux to the atmosphere. However, we did not observe any of the mentioned changes. This points to a potentially active methane biofilter even under anoxic conditions. Notably, potential methane oxidation rates and relative MOB abundance were consistently highest below the oxycline, and both drastically decreased by the onset of water column mixing and reoxygenation in autumn.

### MOB responsible for anaerobic methane oxidation

The high abundance of putatively aerobic MOB in the anoxic water column and their potential activity is an emerging paradox in a variety of aquatic ecosystems. Anaerobic methane removal is commonly attributed to slow-growing methanotrophic archaea in the sediment, where total microbial biomass is high and substrate residence time can be long. Archaea can also be involved in anaerobic methane removal in the water column. For example, in the Black Sea, methanotrophic archaea are responsible for anaerobic methane removal in the suboxic zone (Schubert *et al*., 2006). The relative abundance of methanotrophic archaea in the water column of marine Lake Grevelingen was highest in July with 2 % (of archaeal 16S rRNA reads) at 23 m and 1.8 % at 40 m and mainly belonged to the family of *Methanoperedenaceae*. However, their depth distribution did not follow any coherent pattern and based on metagenomic analysis, in September 2020, the total abundance of archaea in the water column was 50-100 times lower than the abundance of bacteria (based on *phyloflash* analysis of the metagenome, data not shown). Moreover, the slow-growing archaea might struggle to establish a population in shallow ecosystems where the water column is fully mixed in spring and autumn and changes in water column chemistry are fast (Su *et al*., 2023). Therefore, we estimate the overall contribution of methanotrophic archaea to the anaerobic removal of methane in the water column to be low. It is more plausible that the dominant methanotrophic bacteria use alternative electron acceptors during oxygen limitation (Steinsdóttir, *et al*., 2022a). The methanotrophic community in the suboxic zone in July consisted of *MP* and *MMG2*. In the euxinic zone below that, the methanotrophic community was dominated by *MP* alone. The shift in the MOB community towards *MP* and *MMG2* in the anoxic zone may be explained by their versatile metabolism and adaptation potential. Methanotrophic metagenome-assembled genomes (MAGs) previously retrieved from Lake Grevelingen implied the genomic signature of low-oxygen adaptation; a *MP* MAG contained the high-affinity bd-oxidase, which indicates the ability to scavenge oxygen at very low concentrations, and in a MAG attributed to MMG2, the potential use of nitrate or metal-oxides as alternative electron acceptors was described (Venetz et al., 2023). Recent studies further indicate the high potential for anaerobic methane removal by MOB in the water column of marine ecosystems (Thamdrup *et al*., 2019; Steinsdóttir, *et al*., 2022b), and laboratory incubations indicate the possibility of methane oxidation with iron-oxides by methanotrophic bacteria (Zheng *et al*., 2020; Li *et al*., 2021). Suboxic zones harbour electron acceptors other than oxygen, such as metal-oxides or nitrate (Murray et al., 1999). Although cryptic cycling could arguably obscure the supply of oxygen and alternative electron acceptors, nitrate is depleted in the suboxic zone (except for Aug/Sept), but iron oxides are potentially available (Żygadłowska, *et al*., 2023b). Furthermore, recent studies suggest that external electron transfer to or with dissolved organic matter (DOM) might increase the methane oxidation capacity in wetland and brackish sediments and even in the water column of humic bog lakes (Valenzuela *et al*., 2019; Olmsted *et al*., 2023; Pelsma *et al*., 2023). Considering the high sedimentation in the Scharendijke basin (Egger *et al*., 2016) and eutrophic state of the lake, external electron transfer linked to DOM and metal-oxides might be an additional mechanism supporting anaerobic methane oxidation especially in early summer (July). Therefore, methane oxidation with alternative electron acceptors would potentially enable the removal of methane in anoxic water.

### Seasonal dynamics of water-air methane fluxes

Calculations based on benthic methane flux measurements and the distribution of methane in the water column show that ebullition is a major contributor to the methane flux all year long (Żygadłowska, e*t al*., 2023b). These bubbles can bypass the microbial methane filter directly, or dissolve while travelling upwards through the water and can considerably lower the methane filtering efficiency throughout the year (Żygadłowska, *et al*., 2023b). In contrast, the *in situ* diffusive water-air fluxes of methane followed a clear seasonal pattern (Figure 1A). While the fluxes increased from March to April, the fluxes were low in June when the water column was stratified and methane concentrations had increased in the bottom waters. Together with the high abundance of MOB in the water and high potential methane oxidation rates (Figure 2A; Figure 1C), this suggests that the active microbial methane filter during summer stratification mitigated most of the diffusive methane emissions to the atmosphere. However, in September, the diffusive fluxes of methane were higher despite the abundance of methanotrophs. While the methane water-air fluxes were lower during summer stratification (0.12 – 0.22 mmol m^-2^ d^-1^), the fluxes started to increase towards the end of the stratification period (Figure 1A). Although the methanotrophic abundance in the anoxic water column was high at the end of August (up to 10 %), methane still bypassed the MOB-filter, and methane fluxes to the atmosphere increased at the end of August (0.96 mmol m^-2^ d^-1^) and in September (1.15 mmol m^-2^ d^-1^). While in the preceding months, the methane concentrations right below the oxycline never exceeded 0.5 μmol L^-1^, methane concentrations in the oxycline at 50 % oxygen saturation were 1.6 μmol L^-1^ in September (Supplementary Data “water_column_chemistry”). Interestingly, the methanotrophic community did not decrease significantly, which indicates that the seasonal dynamics in water column chemistry affected methane oxidation rates more than abundance or community structure (Kaupper *et al*., 2020). We suggest a combination of reasons that contribute to increased methane fluxes to the atmosphere from the end of August onwards, despite the high relative the abundance of methanotrophs: 1) the weakening of stratification increased the turbulent flux, which resulted in less time for microbial oxidation 2) nitrite accumulation inhibited methanotrophic activity and linked to that 3) a shift in microbial community structure induced a change in the interactions between the methanotrophic bacteria and other members of the microbial community. The weakening of the water column methane biofilter coincided with the weakening of summer stratification. An intrusion of oxygen through a lateral influx of oxygenated water (Hagens *et al*., 2015; van Haren, 2019) and a weakening of the stratification due to warming could increase the downward flux of oxygen and enhance the upward methane flux (Żygadłowska, *et al*., 2023b). The high abundance of M1B in the anoxic water might be the result of oxygen intrusion prior to sampling at the end of August. The MAG associated with M1B did not reveal any genomic indication for the oxidation of methane during oxygen limitation: neither high-affinity oxidases were found, nor key genes for the utilization of alternative electron acceptors (Venetz *et al*., 2023). Therefore, we suggest that cryptic oxygen intrusions between July and August could have provoked a shift away from MOB adapted to oxygen limitation. This may have led to a temporary weakening of the methane oxidation filter once the introduced oxygen was depleted. In the anoxic water, *Methyloprofundus* and *MMG2* were similarly abundant in August, while *Methyloprofundus* dominated in July. This shift further indicates that intrusions of oxygen-rich water could have affected the community in the anoxic water as well. In September, the methane oxidation rates were much lower compared to the previous month. Notably, at the end of August, nitrite accumulated in the suboxic zone (Żygadłowska, *et al*., 2023b), which could be caused by oxygen inhibition of nitrite oxidation (Sun *et al*., 2021). The NMDS plot based on Bray-Curtis distance shows that while oxygen and methane were most predictive of the bacterial community composition(R^2^ = 0.86), nitrite concentrations could partially explain a shift in microbial community structure (Figure 2B, Supplementary Data “NMDS_output”). It is known that nitrite can inhibit methane oxidation, especially in communities not adapted to nitrite and in some cases, this inhibition appears to be even irreversible (King Gary M. and Schnell Sylvia, 1994; Dunfield P and Knowles R, 1995). The abundance of nitrifiers was very low in the microbial community and nitrite concentrations never exceeded 2 μmol L^-1^in the proceeding months below the oxycline (Supplementary Data “water_column_chemistry”). Therefore, the methanotrophic community was likely poorly adapted to nitrite, and nitrite accumulation might have irreversibly inhibited methanotrophic activity in and resulted in low methane oxidation rates in September (Figure 1B). This could have been accompanied by a cascading effect within the plankton community due to the above-mentioned oxygenation event. For example, it has been shown that methane oxidation can be positively or negatively influenced by the total microbial community (Gilbert *et al*., 2012), through nutrient competition and cross-feeding, and either the production or removal of toxic compounds by heterotrophic bacteria (Ho *et al*., 2014; Krause *et al*., 2017; Veraart *et al*., 2018). Therefore, factors accompanying the oxygenation event could have inhibited methane oxidation activity of *MP* and *MMG2* in the anoxic water. All findings together suggest that a weaker stratification, and increased oxygen supply together with potential nitrite accumulation and associated shifts in the microbial community might have resulted in the malfunctioning of the microbial methane filter and resulted in even higher diffusive methane fluxes to the atmosphere.

## Conclusion

We conclude that the efficiency of the microbial methane filter in the water column of a shallow marine basin is strongly influenced by seasonal water column dynamics. The microbial community consists of potentially metabolic versatile *Methylomonadaceae* and counteracts high benthic methane fluxes during summer stratification. This is strikingly reflected in high water column methane oxidation potential and lower *in situ* diffusive water-air methane fluxes. Interestingly, a high abundance of methane-oxidizing bacteria and a high potential for aerobic methane oxidation were even found in the anoxic water. We suggest that methanotrophic bacteria dominate anoxic methane removal in the water column due to the availability of iron-oxides and/or external electron transfer in the suboxic zone. However, methane increasingly bypasses the biofilter as stratification weakens and the contribution of the ebullitive flux from the sediment directly to the atmosphere is high. Hence, notwithstanding the large capacity of the microbial methane filter to oxidize methane under varying redox conditions, water column mixing can decrease the methane filtering efficiency: either directly by higher vertical turbulent flux of dissolved methane or by altering other potential drivers for methane oxidation such as nutrient limitation, changes in microbial community structure or inhibition. This contradicts the assumption that mixing events increase methane oxidation by re-oxygenating the system. Our study demonstrates that the oxygenation state of the water column is not the ultimate factor that determines the functioning of the microbial methane filter. To accurately predict methane emissions from seasonally dynamic eutrophic coastal basins, a more complex network of drivers should be considered.

## Acknowledgements

We are grateful for the excellent support of the captain and crew of the R/V Navicula during the sampling campaigns. Furthermore, we thank Koen A.J. Pelsma and Damian L. Arevalo Martinez for sharing their expertise on floating chamber gas flux measurements. For the dedicated planning and construction of the floating chamber for the flux measurements we thank Arjan de Kleine and the team of the Techno Centrum at Radboud University, Nijmegen (NL). We also want to thank the master students Selien Op den Camp, Andreea Marin and Sam Hoogars for their enthusiastic assistance in the lab. The financing was provided by ERC Synergy grant MARIX (8540088), the program of the Netherlands Earth System Science Center (NESSC 024002001) and Marie Skłodowska-Curie Co-fund (No 847504). NWO VI.Veni.222.332.

## Notes

### Competing Interest Statement

The authors have declared no competing interest.

### Summary of Updates

Matched online abstract with abstract on PDF file

