## Supplementary Information for "Seasonal dynamics of the microbial methane filter in the water column of a eutrophic coastal basin"

#### Supplementary methods

The methane concentrations in the subsamples ($c_a$) were calculated with Henry's law. Measured headspace concentrations ($c_g$) were multiplied by the Henry solubility coefficient $H^{cc}$ (Sander, 2015):

$$c_a = H^{cc} c_g \qquad [1]$$

Where the Henry solubility coefficient is defined as follows: $H^{cc} = H^{cp} RT = \beta \frac{1}{RT^{STP}} RT$ [2]

$H^{cp}$: Henry solubility coefficient (defined as $c_a/p$)
R: ideal gas constant (8.314 J $mol^{-1}$ $K^{-1}$)
T: temperature (294.14 K)
$T^{STP}$: the standard temperature for the Bunsen coefficient (273.15 K)
$\beta$: Bunsen coefficient (taking into account salinity and temperature)

We accounted for the changes in solubility due to salinity and temperature in the calculation of the Bunsen coefficients (Weiss 1970):

$$ln\beta = A_1 + A_2\left(\frac{100}{T}\right) + A_3 ln\left(\frac{T}{100}\right) + S\left[B_1 + B_2\left(\frac{T}{100}\right) + B_3\left(\frac{T}{100}\right)^2\right] \qquad [3]$$

$A_{1\text{-}3}$, $B_{1\text{-}3}$: Bunsen constants, specific for gas
T: Temperature (294.15 K)
S: Salinity (30 ‰)
Next, Bunsen coefficients for $CH_4$, $CO_2$ and $O_2$ were calculated according to the specific constants for each gas (Weiss, 1970, 1974; Yamamoto *et al.*, 1976).

### Supplementary figures

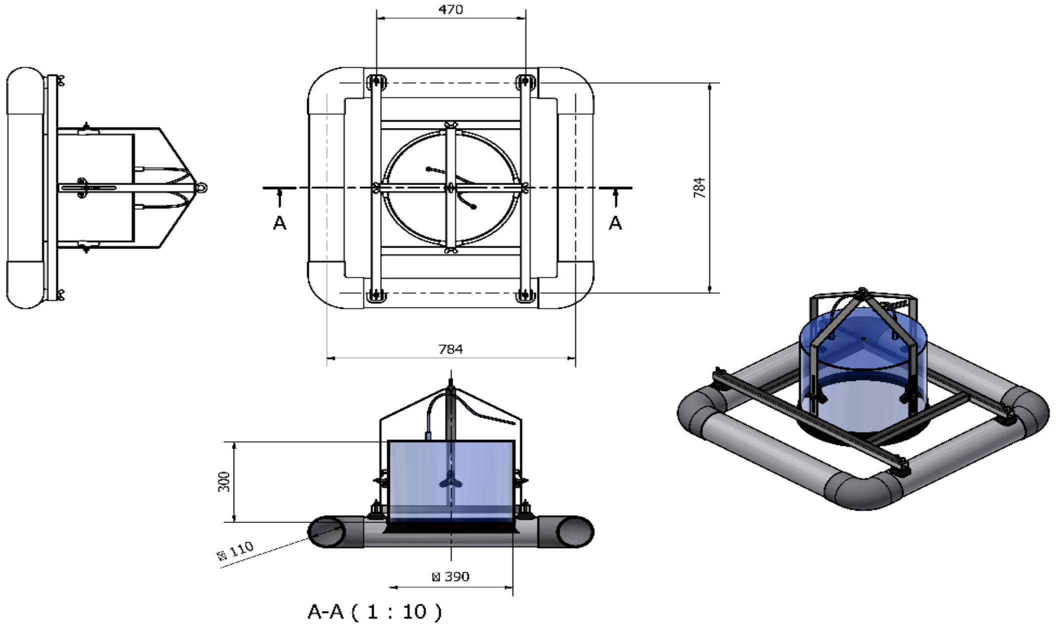


**Figure S1:** *Construction plan of the floating chamber used for in situ flux measurements. Measurements are given in cm. Construction was planned and made by the TechnoCentrum at Radboud University, Nijmegen (NL).*

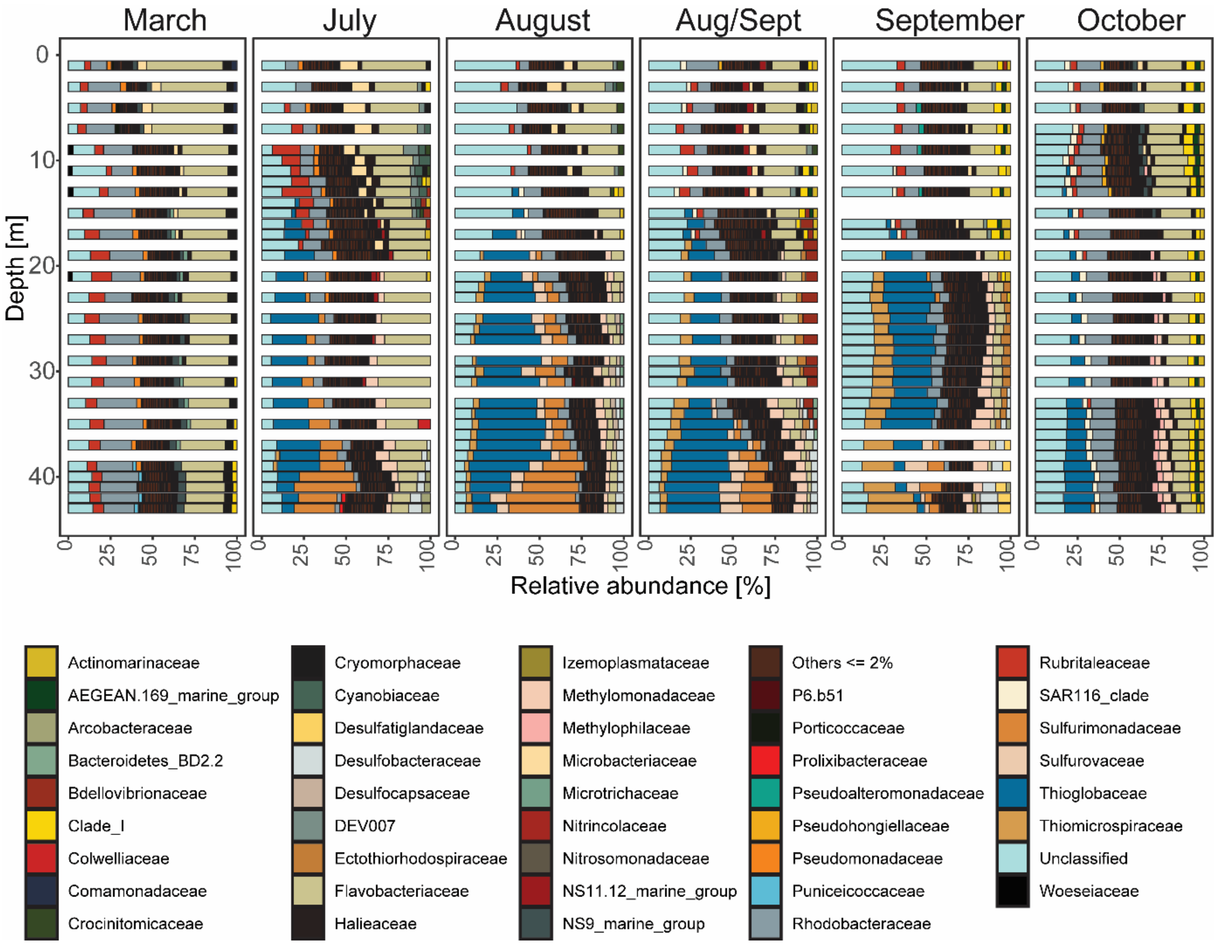


**Figure S2:** *Depth profiles of relative abundances of bacterial family counts retrieved by 16S rRNA illumina sequencing***.**
